# CellDrift: Inferring Perturbation Responses in Temporally-Sampled Single Cell Data

**DOI:** 10.1101/2022.04.13.488194

**Authors:** Kang Jin, Daniel Schnell, Guangyuan Li, Nathan Salomonis, V. B. Surya Prasath, Rhonda Szczesniak, Bruce J. Aronow

## Abstract

Cells and tissues respond to perturbations in multiple ways that can be sensitively reflected in alterations of gene expression. Current approaches to finding and quantifying the effects of perturbations on cell-level responses over time disregard the temporal consistency of identifiable gene programs. To leverage the occurrence of these patterns for perturbation analyses, we developed CellDrift (https://github.com/KANG-BIOINFO/CellDrift), a generalized linear model-based functional data analysis method capable of identifying covarying temporal patterns of various cell types in response to perturbations. As compared to several other approaches, CellDrift demonstrated superior performance in the identification of temporally varied perturbation patterns and the ability to impute missing time points. We applied CellDrift to multiple longitudinal datasets, including COVID-19 disease progression and gastrointestinal tract development, and demonstrated its ability to identify specific gene programs associated with sequential biological processes, trajectories, and outcomes.

## Introduction

Single-cell transcriptomics sequencing (scRNA-seq) has revolutionized discoveries in complex biological systems by identifying a wide variety of cell populations in a high resolution [1–3]. Researchers have applied the technology in experiments with perturbation settings such as diseases [4,5], treatments [6,7], genetic mutations [7,8], organ differentiation [9,10] and more to explore transcriptional profiles across control and various biochemical states. Additionally, large-scale experimental methods, such as CROP-seq [11] and Perturb-seq [12], have been developed for perturbation screening, which provides an abundance of information about biological states.

However, the response to perturbation can vary over time, which is overlooked in many single cell studies. Nowadays, researchers are increasingly considering the impact of time when designing experiments. For example, the genetic effects of autism risk genes have been studied during the development of the nervous system using brain organoids [13,14]. Additionally, influenza vaccination effects have been evaluated by monitoring immune responses over multiple follow-up periods [15]. Moreover, the impact of infections, such as COVID-19 and HIV, has been studied for patients at varying stages of their illness [16,17]. By having access to single cell profiles over time, researchers can accurately report perturbation effects during treatment procedures, disease progression, and organ development.

There have been both conventional and novel approaches introduced to quantify and disentangle transcriptional changes in scRNA-seq data from perturbation experiments [18] (Table 1). Although traditional methods, such as the Wilcoxon test or t-test, are commonly used in single-cell differential expression analysis, they are not sufficient to resolve batch effect and data sparsity issues [19]. More advanced algorithms, such as MAST [20] and muscat [21], have been developed. However, their flexibility in measuring perturbation effects is still limited, such as the identification of common and cell type specific perturbed genes. In the meanwhile, machine learning approaches and generative models have been developed to solve complicated perturbation data. For example, Augur used cross-validation scores of random forest to prioritize perturbation effects across cell types [22]. scGen and CPA applied autoencoder models to learn perturbation responses in the latent space and predict unseen scenarios [23,24]. Although machine learning approaches are powerful in analyzing high-dimensional data, interpretability in latent spaces remains a significant challenge and more so in temporal variation settings.

**Table 1.** Comparisons of perturbation or temporal evaluation methods in single-cell analysis.

| Method | Algorithm | Feature Space | Temporal Evaluation | Reference |
| --- | --- | --- | --- | --- |
| CPA | Autoencoder + linear model | Neural network latent space | Linear model covariate | [24] |
| MEFISTO | Factor analysis | Factors | Gaussian Process | [25] |
| CellBox | Ordinary differential equation (ODE) | Protein and phenotypes | ODE | [27] |
| scGen | Autoencoder | Neural network latent space | N/A | [23] |
| MAST | Generalized Linear Model | Genes | Manual Comparison | [20] |
| CellDrift | Generalized Linear Model | Genes | Functional Data Analysis | This paper |

In addition to the lack of interpretability of the current methods, another defect is that they were not originally designed to measure perturbation effects in a temporal context. Some temporal approaches, such as MEFISTO [25], introduced a regularized factor analysis-based approach to evaluate temporal patterns in single-cell data. These approaches, however, take a transcriptional profile rather than perturbation effects into account. Other methods, such as compositional perturbation autoencoder (CPA) [26], utilized the combination of linear models and deep-learning approaches to interpreting time impact by adding it as a covariate. However, the autoencoder neural network framework and predicted perturbation effects on latent space prevent users from interpreting gene-level impacts simply. Furthermore, CellBox analyzes the perturbation effects over time using Ordinary Differential Equations (ODE) [27]. However, the performance of the method is limited by computational challenges, such as training efficiency on large-scale data, as well as the sparsity and stochasticity issues in single-cell data.

Generalized linear models (GLM) have been widely used in modeling single-cell transcriptomics data, outperforming linear regression by more accurately and efficiently capturing non-linear relations in the count data through non-Gaussian distributional families formed by link functions [28]. Furthermore, it is more flexible in the modeling of the mean-variance relationship of the count data. Sctransform successfully removed technical effects by introducing cellular sequencing depth as a covariate [29]. Milo captured differential abundance in single-cell perturbation data by adding cell counts as the covariate in a GLM on k-nearest neighbor (KNN) graphs [30].

Functional Data Analysis (FDA) has been widely used in longitudinal data analysis [31,32]. A general form of FDA is the analysis of multiple curves varying over time, where each curve is a sample tracing with a series of time points, which can be characterized as a function. Such data is called functional data. One of the most popular tools in FDA is Functional Principal Component Analysis (FPCA), which identifies the dominant modes of variation of functional data [33]. It has been widely used in disease progression profiling and predictions. For example, FPCA has been used in the trajectory evaluation of cystic fibrosis progression [34] and monitoring of glucose levels in hyperglycemic patients [35]. We utilized the flexibility of the FDA to identify temporal perturbation patterns.

To address the aforementioned issues, we developed CellDrift, a generalized linear model-based functional data analysis model to disentangle temporal patterns in perturbation responses in single-cell RNA-seq data. CellDrift first captures cell type specific perturbation effects by adding an interaction term in the GLM and then utilizes predicted coefficients to calculate contrast coefficients, which represent perturbation effects in our study. Concatenated contrast coefficients over time are defined as functions and Fuzzy C-mean clustering is used to identify temporal patterns, accompanied by FPCA to find major components that account for the most temporal variance. We benchmarked CellDrift with multiple functional clustering methods with statistical results from differential expression approaches, such as Wilcoxon and t-test, and CellDrift achieved superior performance in the identification of temporal patterns and imputation of perturbation effects. We applied CellDrift in COVID-19 single cell data and gut development atlas and identified temporal patterns and functional principal components associated with varying immune responses and gut organ morphogenesis. The CellDrift package is available on https://github.com/KANG-BIOINFO/CellDrift.

## Methods

CellDrift takes the input of multiple single-cell RNA-Seq UMI count matrices across diverse captures (batches, *b*), conditions (perturbations, *p*) and time points (*t*). As the main goal of the algorithm aims to disentangle the major effects of different cell types (*c*) and perturbations (*p*), we utilized the Generalized Linear Model (GLM) due to its superior flexibility and interpretability to model the raw count data in a probabilistic manner. The derived contrast coefficients associated with each gene, cell type and condition after implementing the GLM model across time points are used for functional data analysis to identify temporal patterns of perturbation responses.

### Perturbation coefficient model

To begin, we introduce a model and notation for a single time point. We model the raw count of single-cell data *γ*_*ng*_ for cell *n* and gene *g* using a generalized linear model with a negative binomial (NB) distribution. *Z*_*n*_ and *x*_*n*_ represent cell type and perturbation group of cell *n*, which are *c* and *p* here:

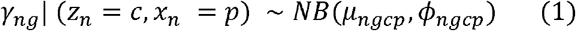

where *µ*_*ngcp*_ and *µ*_*ngcp*_ represent the mean and inverse dispersion of the NB distribution for cell *n* and gene *g*. In real cases, *Z*_*n*_ and *x*_*n*_ are user-defined cell type and perturbation group for cell *n*, such as CD4+ T cell and Drug A treatment.

We use a log link function (*ln*) for *µ*_*ngcp*_ to disentangle *η*_*ngcp*_ with a linear model with cell type coefficients *β*_*gc*_ and perturbation coefficient *β*_*gp*_ for each gene *g*:

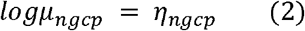

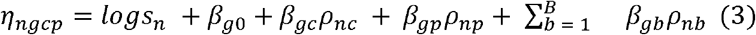

where *ρ*_*nc*_ and *ρ*_*np*_ represent whether cell *n* belongs to cell type *c* and perturbation group *p*, as represented by one-hot matrices. The intercept *β*_*g*0_ represents the base expression level of gene *g*. In addition, we account for library size *S*_*n*_ and batch effects *β*_*gb*_ in the model, and *ρ*_*np*_ is the one-hot matrix for cell *n* and batch *b* (e.g., donor, sequencing platform). *B* is the total number of batch types. Batch types are incorporated into the model as fixed effects. For simplicity, we omit the *b* (batch) subscript in the mean count (*µ*_*ngcp*_) and linear predictor symbols (*η*_*ngcp*_).

In equation (3), cell type effect and perturbation effect are independent covariates. In real cases, however, different cell types usually have distinct responses towards the same perturbation. Thus, we add an interaction term *β*_*gcp*_ for cell type and perturbation covariates, representing cell type-specific perturbation effects. In contrast, *β*_*gp*_, represent common perturbation effects across cell types:

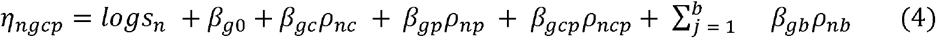

where *ρ*_*ncp*_ is a one-hot matrix of cell type and perturbation group, indicating whether cell *n* belongs to cell type *c* and perturbation group *p* at the same time.

By adding the interaction term *β*_*gcp*_, we create a contrast model to determine whether or not perturbation effects vary across cell types. To justify the performance improvement of models for some genes by adding the interaction term, we applied the likelihood ratio test (LRT) for the reference and contrast models with the null hypothesis (*H*_0_) that the contrast model doesn’t significantly perform better than the reference model (Figure S1). Using a cutoff (FDR-adjusted *p* values < 0.05) for LRT tests, we then identify genes with cell-type-specific perturbation effects.

### Contrast coefficients

Both major effects (*β*_*gc*_ and *β*_*gp*_) and interaction coefficients (*β*_*gcp*_) are predicted using GLM after fitting the single cell data. Then we use Estimated Marginal Means (EMM) [36] to retrieve pairwise contrast coefficients *Δβ*_*gcp*_ based on predicted *β*_*gp*_ and *β*_*gcp*_, which are used to measure the difference between the perturbed state and baseline in specific cell types. Briefly, they represent the coefficients of cell type *c* (perturbation group *p*) minus cell type *c* (control). Contrast coefficients are the basic representation of perturbation effects in this study. They are also the input data for FDA (Figure 1).

**Figure 1.**
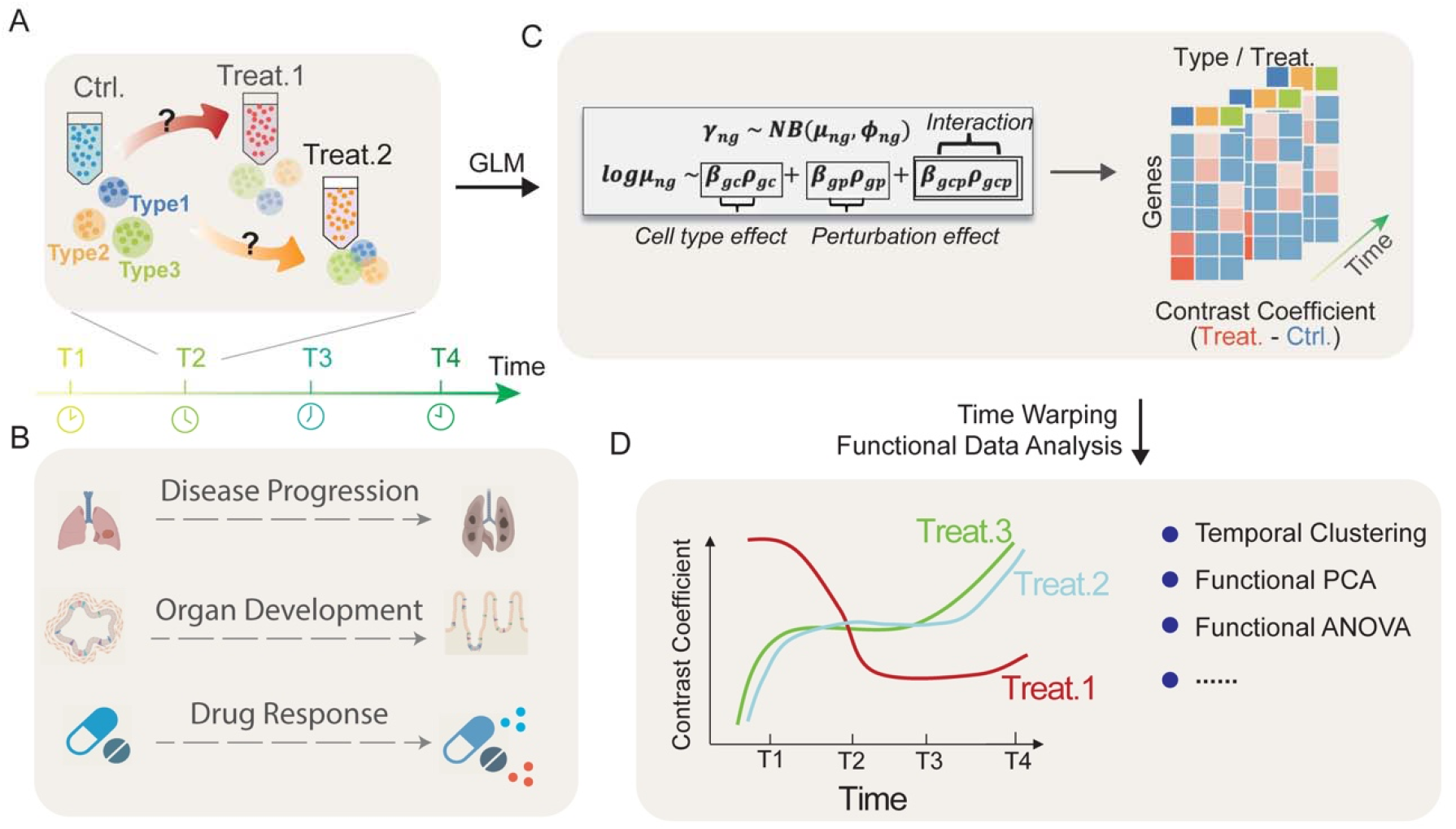
Workflow of CellDrift. (A) An example of a perturbational single cell experiment with multiple time points. (B) Real scenarios of single cell experiments with varying perturbation effects over time. (C) Generalized linear model (GLM) with the interaction of cell-type-perturbation applied separately at each time point and contrast coefficients are derived as the representation of perturbation responses. For simplicity, we omit the library size and batch effect in the linear function. More details can be found in the Methods. (D) Contrast coefficients are used as input for various applications in functional data analysis, including temporal pattern identification, functional PCA and one-way functional ANOVA.

In line with the standard strategy implemented in R package emmeans [36], we first get pooled standard errors for pairwise tests of estimates from the generalized linear model. *Z*-scores are derived by dividing means by the standard errors. Using these scores, we calculate *p* values using normal approximation, which for a two-tailed test is 2 ×the probability of *z*-scores on the negative scale. In CellDrift, *Z*-scores are used as the final values of contrast coefficients.

### Representation of functional data using contrast coefficients

Functional data analysis (FDA) is a popular framework for longitudinal data analysis, whose general form is the analysis of multiple curves over time. Each curve is a sample with a series of time points, which is commonly referred to as a function. As an example, varying glucose levels over time for a patient can be described as a function.

In our study, in addition to cell types and perturbation groups in single-cell perturbation data, we added another dimension of complexity, time covariate *t*, into our model, which is continuous and usually sparse covariate in single-cell experiments. Real-life examples of time covariates include the elapsed time since disease onset, drug treatment, genetic knockouts, and other cases.

In our model, we estimate contrast coefficients *Δβ*_*gcp*_ for genes across cell types and perturbation groups at each time point *t* from the time series {1,2, …, *T*}. Then for each combination of cell type *c* and perturbation group *p*, we get a series of *Δβ*_*gcp*_ across available time points, represented as 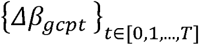. Each series 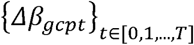 denotes perturbation coefficients of gene *g* across time points for the selected cell type and perturbation group, which is the representation of a function or a sample in the following functional data analysis framework.

### Temporal Pattern Identification

There are two general input formats for FDA in our context:

1. Functional data of various genes in a fixed cell type and perturbation group:

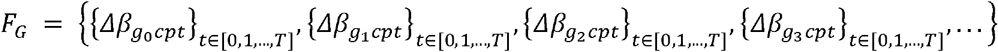
2. Functional data of various cell types and perturbation combinations for a specific gene:

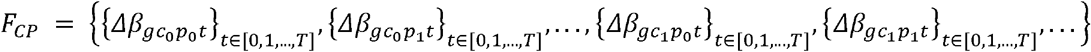

where functional data are denoted as a set of functions.

Our goal is to identify genes (or cell type-perturbation combinations) that show similar dynamic changes in perturbation responses over time, which we refer to as temporal patterns. To find such patterns, we used functional clustering algorithms such as KMeans, Fuzzy C-means, and EMCluster [37,38]. Based on benchmark results, we utilized the Fuzzy C-means functional clustering algorithm to identify temporal patterns of perturbation effects [39]. For example, when clustering on *F*_*G*_, the resultant cluster can be interpreted as a group of genes with a similar pattern of perturbation response over time. L2 distance of functional data was used to iteratively update clustering results. Different from classical K-Means clustering, Fuzzy C-means clustering uses weighted square errors in the objective function, where the algorithm iteratively updates the probability of a sample being a member of a cluster (supplementary materials).

### Decomposing temporal complexity with FPCA

In real cases, it is easy to see thousands or even more samples (genes or cell type-perturbation combinations) in the input data. To decompose the complexity of the data on the time scale, we implemented Functional Principal Component Analysis (FPCA) and extracted top functional principal components (PCs) that explained most temporal variance. To resolve the sparsity issue of functional data due to the lack of time points, we used Principal Analysis by Conditional Estimation (PACE), which yields covariance and mean functions, eigenfunctions and principal component scores for both functional data and its derivatives with respect to time [40,41]. Notably, the covariance of curves in PACE is smoothed, resulting in smoothed temporal curves in the output (supplementary materials). In conclusion, PACE can not only reduce temporal complexity but also generate smooth curves.

### ANOVA Test for Differential Temporal Perturbation Effects

We implemented a one-way functional ANOVA to identify genes with differential patterns across perturbation groups [42]. Let *P*_1_, *P*_2_, …, be functions from different lists of perturbation groups. We define 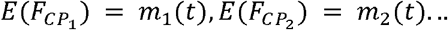, The null hypothesis is:

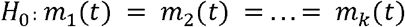

More details can be found in the supplementary materials.

### Time Warping

In some situations, the sampling time points can be variable for different perturbations. For example, single cell data of COVID-19 and sepsis patients were retrieved from different time scales in the same study [43]. We used the dynamic time warping (DTW) algorithm to align different time series. It aligns two time series with different lengths by comparing the similarity or calculating the distance between them [44]. It is necessary when samples in different perturbation groups are collected from unaligned time points. Application of DTW could make temporal curves comparable. We first choose a reference curve and then find the matching indices between query and reference curves. Values of time points on query curves are projected onto the matched time points in the reference curve, making temporal data from different curves comparable.

### Perturbation Data Simulation

Simulation data with varying batch effects and differential expression sizes were generated from *Splatter* [45]. To mimic real single-cell data, we extracted initialization parameters from a CD14+ Monocytes of an interferon-stimulated PBMC dataset [46]. We define 3000 features and 1000 cells with 2 cell groups (Group 1,2) and 3 batches (Batch 1,2,3). To investigate the influence of batch effects, we selected varying batch effect sizes (*batch. facboc*)from [0.02, 0.1, 0.4, 0.7] for 3 batches. *batch. facScale* is controlled as 0.1. The proportion of differential expression (DE) genes compared with baseline (*de*. *prob*) is controlled as 50% for all three batches with shared DE genes, among which 50% are downregulated genes. To simulate prominent batch effects for benchmark tasks, we intentionally introduced imbalance in the largest batch, with 600 cells in Batch 1 and 200 cells in Batch 2 and 3 (Figure 2).

**Figure 2.**
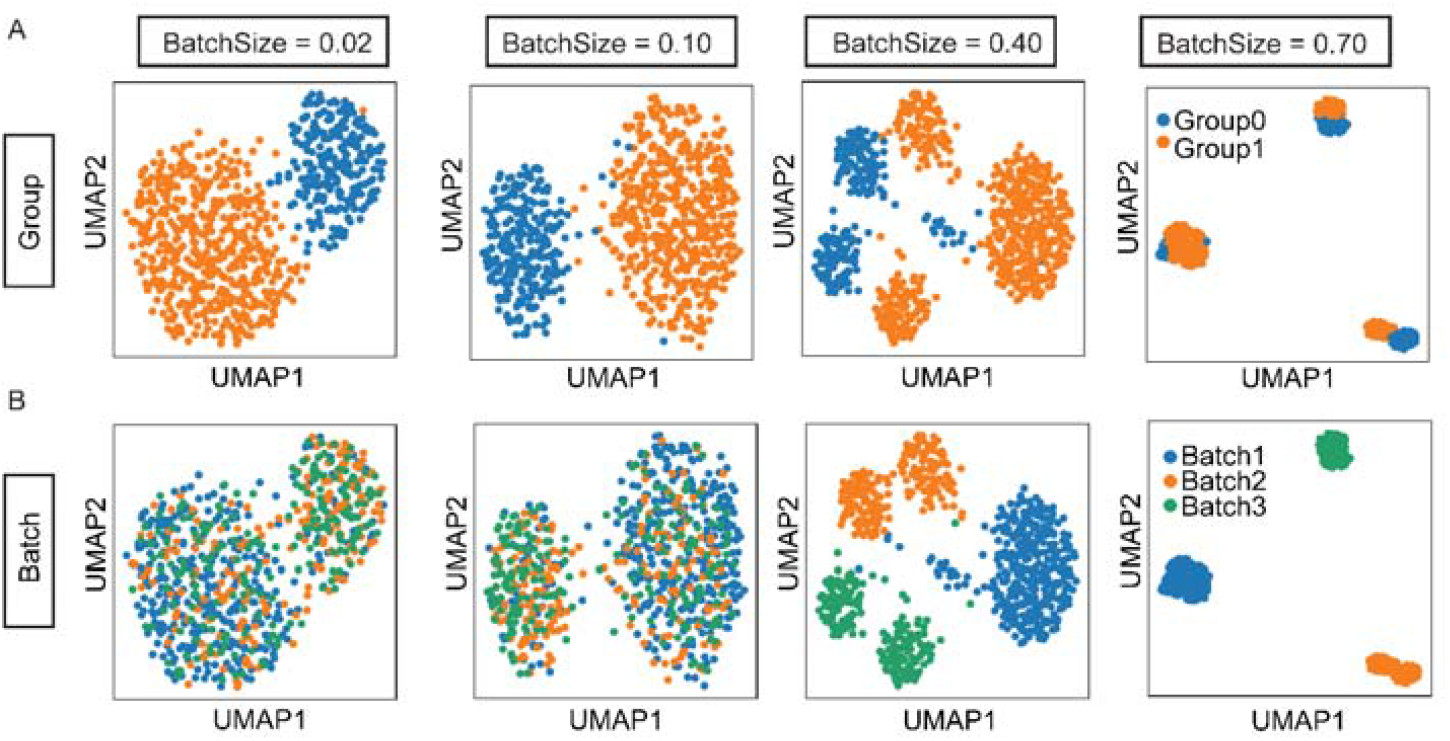
Splatter simulated data for the benchmark of CellDrift generalized linear model. (A-B) UMAPs of simulated data with different batch effect sizes range from 0.02 to 0.7. Batch (A) and biological cell group (B) annotations are shown.

Additionally, to interrogate the influence of differential expression size, the differential expression parameter *de*. *prob* was defined with a series of values ranging in [0.05, 0.2, 0.5, 0.8], representing the proportion of DE genes compared with the baseline expression in the simulated data. To clarify, this parameter doesn’t denote the DE gene proportion between Group1 and Group2. In this benchmark experiment, simulated data contains 600 cells and only one batch.

To define the ground truth of DE genes between Group 1 and Group 2, we divided genes from simulated data into two categories: (i) unperturbed (negative)s genes with differential expression factors in both Group1 and Group2 as 1; (ii) perturbed (positive) genes with absolute difference of between Group 1 and Group 2 in the top 75 percentile.

### Temporal Data Simulation

Inspired by MEFISTO [25], simulation data were derived from a generative model by varying the number of time points per group in a [0, 1] interval, the rate of missing time points, the noise levels and sequencing depth (Figure 3). The default base mean of simulated genes is 2. Default coefficient parameters for cell type, perturbation and the interaction effects are 0, 0.3 and 2, and corresponding scale parameters are 0.2, 0.2, 0.2, respectively. We defined 3 linear temporal patterns in our study and cell type-perturbation interaction coefficient is a time-dependent parameter. By default, are -1, 0, 1 for negative-correlated, time-insensitive and positive-correlated temporal patterns. Corresponding intercepts are 1, 1, 0. We also defined 3 non-linear temporal patterns where (Figure 3E), and default are -4, 0, 4, representing various shapes of non-linear temporal patterns. We also defined as [-8, 0, 8] and [-2, 0, 2] to represent different temporal curve shapes in the benchmark. In each parameter setting, 100 cells in each cell type and perturbation group per time point, and 20 genes were simulated for each temporal pattern. 12 replicates were generated for each parameter setting.

**Figure 3.**
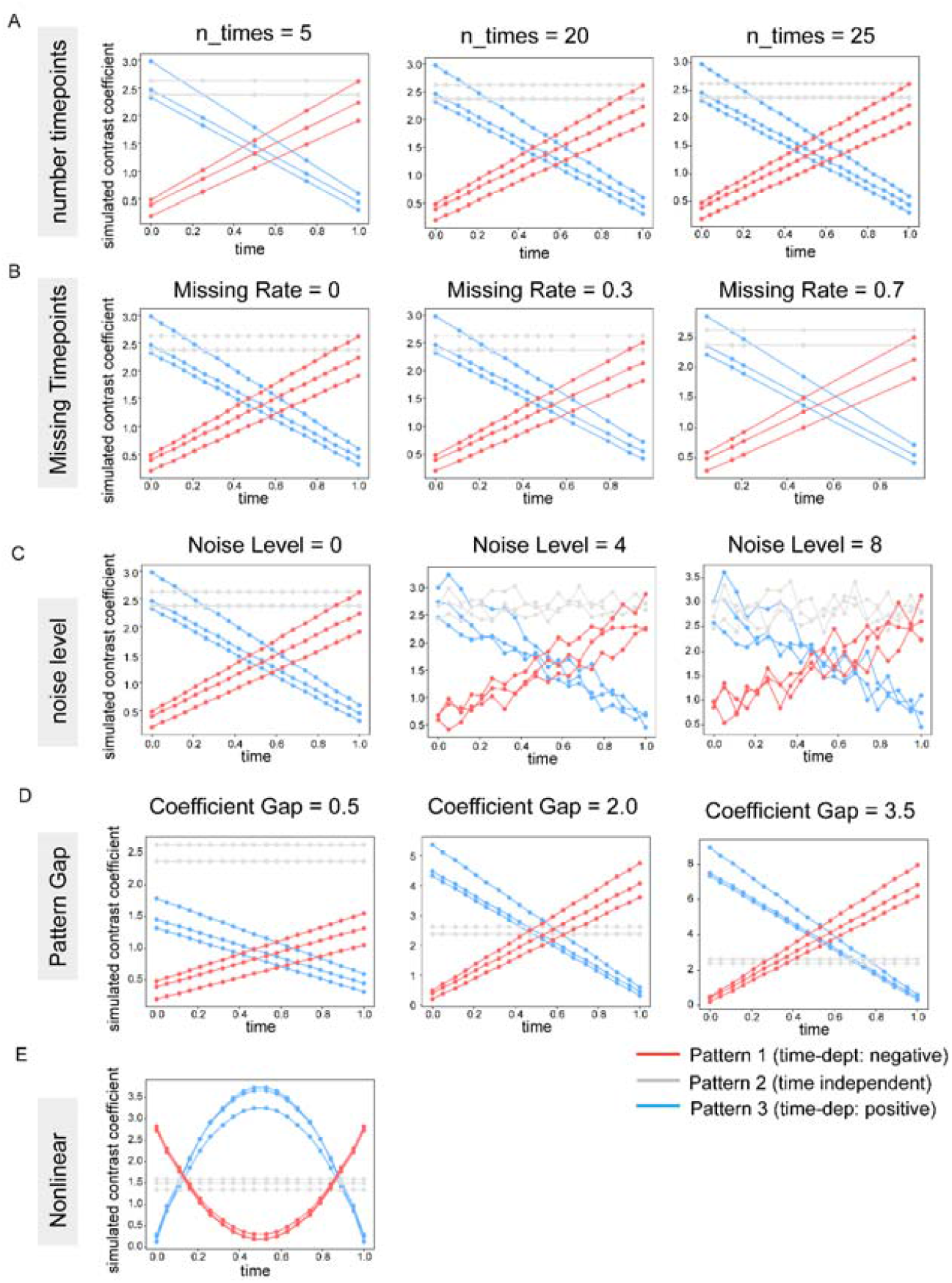
Simulated data for the temporal pattern identification. (A-E) Simulated data with 3 temporal patterns of contrast coefficients, including negatively-correlated temporal pattern (blue), time-insensitive pattern (gray) and positively-correlated time pattern (red). Varying parameters were applied in the simulation algorithm, including the number of time points (A), ratio of missing time points (B), noise level (C), coefficient gaps between time patterns (D) and non-linear time patterns (E). Three random genes in each temporal pattern are shown on the plot.

### Benchmark Criteria

In this study, we benchmarked two parts of the algorithm, including perturbed gene prediction using the generalized linear model and temporal pattern identification using functional data analysis.

As we mentioned previously, true perturbed genes and unperturbed genes were defined in the Splatter simulation data as positive and negative data. We used the true positive rate (TPR or sensitivity) and the false discovery rate (FDR) as metrics to evaluate the sensitivity and false discoveries of GLM in identifying perturbed genes.

In the benchmark experiment of temporal pattern identification, three temporal patterns were defined in the simulation data. Then we used the adjusted rand index (ARI) to measure the similarity between estimated functional clusters *C*_*E*_ with true temporal patterns *C*_*T*_ :

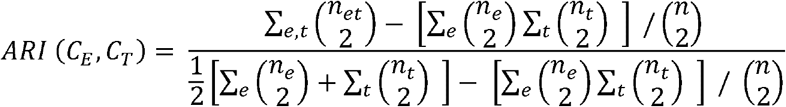

where *n* is the total number of genes; *n*_*e*_ and *n*_*t*_ are the number of genes in each estimated cluster *e* and in the true cluster *t*, respectively; and *n*_*et*_ is the number of genes shared by estimated cluster *e* and true cluster *t*. ARI ranges from 0 to 1, where a larger value indicates more similarity between estimated clusters and true clusters.

Additionally, we utilized FPCA to retrieve smoothed curves of functional data, where we have imputed contrast coefficients for missing time points. To compare imputation performance with real simulated contrast coefficients, we used Pearson correlation to measure the consistency between simulated and estimated contrast coefficients for missing time points in linear temporal patterns.

## Results

### CellDrift Generalized Linear Model improves perturbed gene detection

As a simple application of the GLM model of CellDrift, we first applied it in a interferon-stimulated peripheral blood mononuclear cells (PBMC) single-cell data with a single time point.As expected, genes with the highest positive and negative contrast coefficients are closely linked to inflammatory pathways (Figure 4A, B, Supplementary Table S1) [46].

**Figure 4.**
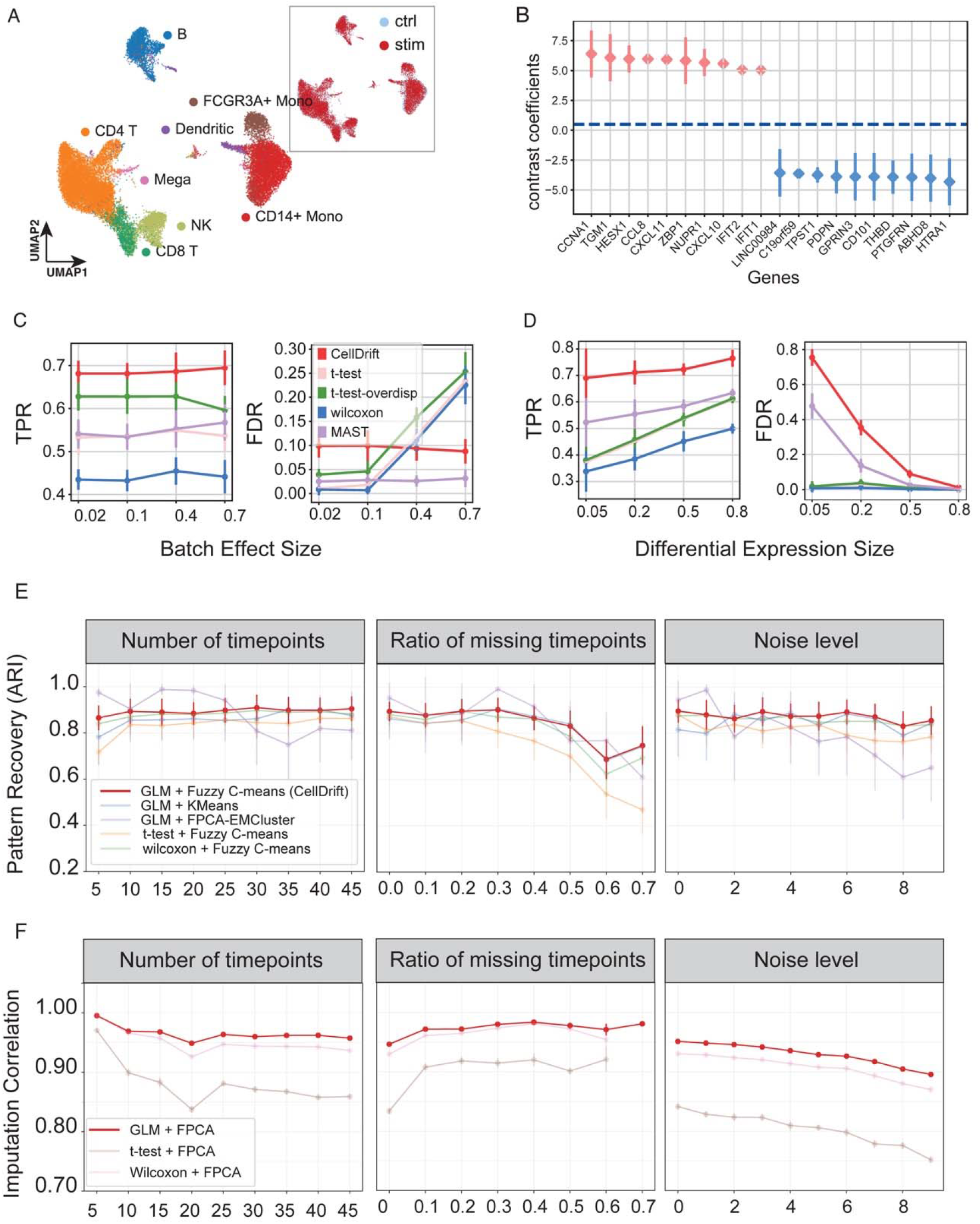
Benchmark of the CellDrift approach. (A-B) A simple application of CellDrift in the interferon-stimulated PBMC single-cell data. (A) UMAP of cell types and treatment groups of PBMC single-cell dataset. Ctrl: control; stim: interferon-stimulated group. (B) CellDrift for 10 genes with the highest positive and negative contrast coefficients for classical monocytes in the interferon-stimulated condition. (C-D) Single time point simulation benchmark of the generalized linear model of CellDrift with other commonly used differential expression approaches. Simulated data with multiple batch effect sizes (C) and differential expression sizes (D) were used for benchmarking. True positive rate (TPR) and false discovery rate (FDR) were used as metrics. The mean and standard deviation derived from 10 replicates for each test are shown in the figure. (E) Benchmark results of temporal pattern recovery for CellDrift strategy (GLM + Fuzzy C-mean) and other approaches. Temporal pattern recovery was evaluated with adjusted rand index (ARI) and measured across varying parameters, including numbers of time points, ratios of missing time points and noise levels. (F) Benchmark results of imputation performance using Pearson correlations between imputed contrast coefficients from FPCA and real simulated perturbation coefficients. The same parameters were used as (E).

Furthermore, we demonstrated the performance of CellDrift in the simulation data. We defined significantly perturbed genes in CellDrift and each benchmarked method as those with FDR-adjusted p values less than 0.05. True positive rates (TPR or sensitivity) and false discovery rates (FDR) were derived by comparing the ground truth (see methods) and estimated results. These evaluation criteria have been widely used in established methods [30]. Compared with other commonly-used differential expression methods, including t-test, wilcoxon test and MAST (supplementary materials), CellDrift achieved improved sensitivity in diverse levels of batch effect sizes and differential expression sizes (Figure 4C,D), which indicates stronger detection power for perturbed genes using CellDrift.

We observed that CellDrift has a higher FDR in experiments with small batch effects (<0.1). However, it has a stable and controlled FDR at larger batch effect sizes (0.4 and 0.7), where higher FDR was observed in other methods, such as t-test and wilcoxon (Figure. 4C). Similarly, MAST has a stable and small FDR of less than 0.05 across different batch effects, showing the best performance of controlling FDR among all methods, which may be due to the removal of technical covariates, such as batch effects, by the hurdle model [20]. However, its TPR is much lower than CellDrift.

CellDrift outperformed other methods with significantly higher TPR across various differential expression sizes (Figure 4D). Meanwhile, we observed a high FDR of CellDrift in experiments with small differential expression sizes (0.05 and 0.2), indicating relatively inferior performance in controlling false discoveries. MAST had similar results. We argue, however, that it is more important to identify as many DE genes in rarely perturbed data as possible than to avoid false discoveries (Figure. 4D). FDR of CellDrift decreased with increasing differential expression sizes, and achieved a low level (<0.15) in large DE sizes (0.5, 0.8).

### Fuzzy C-means clustering and CellDrift contrast coefficients improve temporal pattern identification and imputation performance

We simulated both linear and nonlinear time patterns of gene perturbation effects. The temporal pattern recovery performance of three functional clustering algorithms was examined, including KMeans, Fuzzy C-means, and FPCA-based EMCluster [47] for CellDrift GLM contrast coefficient input. We used Adjusted Rand Index (ARI) as the metric to measure the accuracy of prediction for simulated temporal patterns. The influence of different parameters, including the number of time points, ratio of missing time points and noise levels were measured (Figure 4E, S3).

From the results, we observed that FPCA-based EMCluster achieved higher accuracy in a certain number of time points. Nonetheless, Fuzzy C-means achieved stable and better performance than most other methods at varying numbers of time points, with ARI reaching 0.9. Additionally, the ARI of Fuzzy C-means exceeded 0.8 at ratios of missing time points less than 0.5. ARI decreased at greater ratios of missing time points, but remained at the 0.7 level and outperformed most other clustering algorithms, such as the Wilcoxon test. Note that the ARI of Fuzzy C-mean remained stable at high noise levels with relatively higher accuracy than normal KMeans and other methods (Figure 4E). Furthermore, Fuzzy C-mean has shown improved performance for a variety of non-linear time patterns, pattern coefficient gaps and sequencing depth (Figure S3A). Our findings suggest that Fuzzy C-mean has the most stable and relatively better temporal pattern recognition performance.

We also compared GLM-based contrast coefficients with other statistical scores, such as scores from t-test and Wilcoxon test, as the input for functional data analysis (Figure 4E). T-test scores had inferior performance in most parameter settings. The Wilcoxon score achieved better performance than the t-test, and its performance in various time points is comparable with GLM-based functional clustering. However, its performance in large fractions of missing time points is inferior to GLM-based clustering (Figure 4E).

Additionally, we investigated the imputation performance for missing time points, which is a commonly seen situation in real temporal single cell data. Incorporated in our pipeline, FPCA provides smoothing and interpolation functions. We compared GLM-based input with statistical scores from t-test and Wilcoxon test, where we observed significant improvement in imputation performance using GLM-based input (Figure 4F, S3B).

### CellDrift identified temporal patterns of COVID-19 immune responses

We next demonstrated CellDrift by identifying temporal patterns of immune responses in a large-scale COVID-19 PBMC single cell data. Multiple COVID-19 conditions, including mild, mild-HCW (mild healthcare workers), severe and critical patients, as well as severe influenza and sepsis patients. The six most common cell types were extracted for our analysis. CellDrift was used to analyze temporal trends using days since disease onset for patients in the original study (Figure 5A).

**Figure 5.**
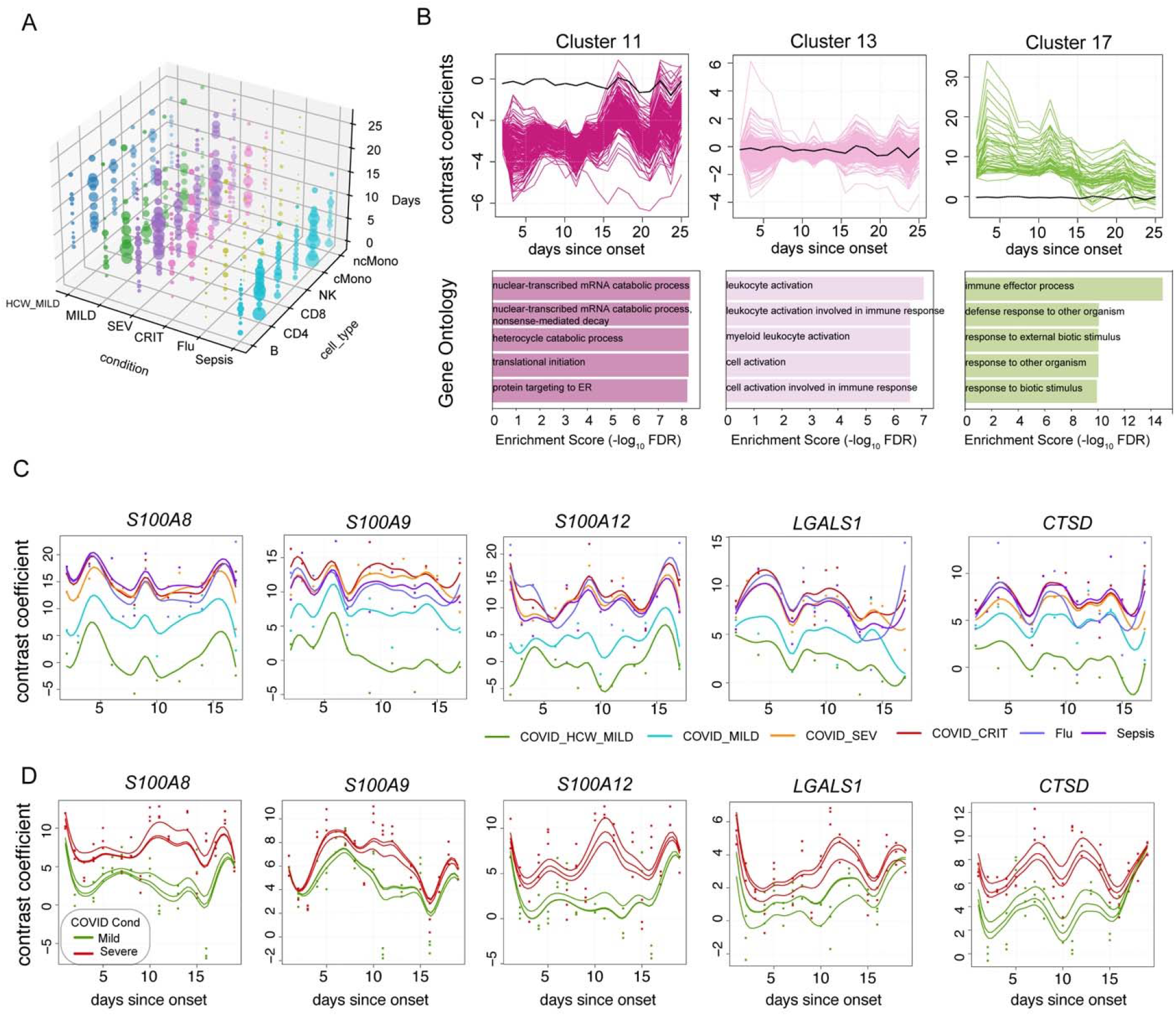
Temporal perturbation effects in COVID-19 atlas. (A) Overview of the number of cells in each cell group of the dataset, which contains 6 disease conditions, 6 selected cell types and time points from day 1 to day 25 since disease onset. The size of dots represents the number of cells. HCW_MILD: healthcare workers with mild COVID-19; MILD, SEV, CRIT: mild, severe and critical COVID-19; CD4, CD8: CD4 T cell, CD8 T cell; cMono, ncMono: classical and non-classical monocyte. (B) Three distinct temporal patterns of contrast coefficients from classical monocytes of severe COVID-19 patients. The top row shows curves of genes with similar contrast coefficients in each cluster over time, and the bottom row shows the gene set enrichment analysis of genes in each temporal cluster. The black line represents the average time curve for all genes, which is the same across three plots. Gene enrichment scores are defined as − *log* _10_ *FDR* adjusted p values of enrichment significance and biological processes from Gene Ontology were selected. (C) Five genes from cluster 17 were prioritized by the one-way functional ANOVA test, which have significantly higher temporal curves in severe conditions than mild symptoms. Contrast coefficients for classical monocytes across disease conditions are shown on smoothed curves computed by FPCA, and time curves were aligned using dynamic time warping. (D) Validation of genes in (C) in another large-scale COVID-19 PBMC data [1]. FPCA smoothed curves for genes in three replicates of mild and severe patients are shown, which display similar temporal patterns as (C).

We first focused on classical monocytes in severe COVID-19 patients and applied CellDrift Fuzzy C-mean for all genes after the feature selection (supplementary Table S2). Genes with similar temporal patterns of perturbation responses clustered together, indicating that several patterns of dynamic changes were occurring upon virus infection during disease progression. The genes responding to perturbations in clusters 11, 13 and 17 showed three distinct temporal patterns, where the contrast coefficients of clusters 11 and 17 showed a positive and negative correlation with time, while cluster 13 showed an insensitive pattern to time (Figure 5B). Based on gene enrichment results, cluster 11 is highly associated with catabolic and biosynthesis processes, while cluster 17 appears to be involved in immune responses, indicating a rapid activation of immune response activities and a suppression of house-keeping activities in the early disease stage (d1 ∼ d15), with a reduced level of such change in later stages (after d15). Additionally, we did functional PCA for all genes and identified 3 eigenfunctions that explained over 99% temporal variance (Figure S4). The first eigenfunction can account for the majority of observed temporal patterns, as shown by the FPC 1 scores in Figure S5. Moreover, genes stratified by FPC1 scores represented positive-correlated, negatively-correlated and insensitive temporal patterns (Figure S6).

After we obtained temporal patterns in severe COVID-19 patients, the next step was to examine whether the patterns vary across multiple perturbation groups, such as mild and severe COVID-19 patients. To achieve it, we applied functional analysis in classical monocytes of multiple disease conditions, where dynamic time warping was used to align multiple time series into a comparable time scale. Next, we applied a one-way functional ANOVA test and calculated ANOVA scores for each gene, representing the consistency of perturbation responses between disease conditions over time. Based on ANOVA results, a number of genes from cluster 17 were identified as severe-prominent genes, including S100A8, S100A9, CTSD and others (Figure 5C). In agreement with our findings, elevated levels of calprotectin (S100A8/S100A9) have been found in severe COVID-19 patients with poor clinical outcomes [48,49]. Apart from severe-prominent genes, we also prioritized mild-prominent genes and condition-irrelevant genes, representing distinct gene programs of temporal perturbation responses across disease conditions (Figure S9).

To validate our discoveries, we also applied CellDrift to data from another large-scale COVID-19 single cell experiment [1]. We observed similar temporal patterns between mild and severe COVID-19 patients as shown in Figure 5C, which shows the reproducibility of CellDrift approach.

### CellDrift discovered differential temporal gene patterns during fetal gut developments

We further implemented CellDrift in a fetal gut cell atlas to identify differential gene programs during organ development [50]. Researchers examined gut development in 3 compartments, including duo-jejunum, ileum, and colon, at 9 time points during development from the embryonic stage (week 6) to the fetal stage (week 11) (Figure S8A). We selected epithelial and mesenchymal cells from duo-jejunum as a reference tissue in order to provide an adequate number of cells to GLM. Differential gene programs during development between the colon (or ileum) and duo-jejunum were generated by GLM across time points (Supplementary Table S3). Notably, the top two eigenfunctions explain more than 99% of the temporal variance of mesenchymal cells between the colon and duo-jejunum, where the first eigenfunction shows reverse temporal patterns during the development (Figure S8B, S9). CellDrift identified 20 temporal clusters (Figure S10), and ranked them by FPC1 scores, in which clusters 11 and 16 had high positive and negative correlations with FPC1 scores (Figure S8C). As a result of gene enrichment, extracellular matrix organization genes are more active in early stages (weeks 7-8) in the duo-jejunum, and then in later stages (weeks 9-10) in the colon, whereas morphogenesis genes are more prominent in distal tissues (colon) at the beginning, and proximal (duo-jejunum) later on. Similarly, the second eigenfunction in the comparison of epithelium cells between colon and duo-jejunum revealed downregulated temporal pattern around early-stage (week 7.9) and upregulated pattern at a later stage (week 9.2) (Figure S8F, G, S9), which were associated with lipid metabolic process and chromosome organization, respectively (Figure S8H, I). As a result of the time lapse of specific gene programs during organ morphogenesis, our findings reveal temporal patterns that appear like waves from proximal to distal compartments throughout the gut development.

## Discussion

In this study, we presented a framework to identify temporal patterns of perturbation responses. As far as we know, CellDrift is the first method to use functional data analysis to evaluate longitudinal perturbation effects in single cell data. Using generalized linear models, we modeled perturbational single cell data and introduced the new concept of cell type-perturbation interaction, which improves the sensitivity of detecting both common and cell type-specific perturbation effects in real-life single cell experiments. As a result of allowing for batch covariates, we address a significant barrier to finding real perturbed genes.

Unlike currently available single-cell methods which either focus on temporal analysis or perturbation investigation, we utilized the flexibility of GLM and functional data analysis to combine these two areas together, and gained insights into evaluating complicated longitudinal perturbation responses. We can use GLM to calculate gene-level perturbation effects instead of latent space features, such as the ones evaluated with CPA and scGen. In many cases, researchers focus on the perturbation effects of specific genes. The COVID-19-induced cytokine storm, as an example, is being studied for temporal perturbation of cytokine genes [6].

In our study, we successfully improved true positive rates in multiple settings of batch effects and perturbation effect size compared with popular methods in differential expression analysis, enabling the capture of more perturbed features. The false discovery rate is insensitive to varying batch sizes, indicating the successful repression of batch effects by CellDrift. Although the false discovery rate was high with small perturbation effect sizes, we argue that enhancing sensitivity is more important than the false discovery rate where genes are rarely perturbed. Notably, although the performance of a generalized linear model with the Fuzzy C-mean was not uniformly superior in identifying temporal patterns, it was the most stable approach and performed well in the majority of benchmark experiments. Similarly, combining the Wilcoxon test with Fuzzy C-mean also delivered good results, but the performance decreased with a higher ratio of missing time points, which is typical in real data. Gaussian Process (GP) has been found to be effective in inferring temporal patterns from single-cell data [25]. FDA was selected over the option of Gaussian Processes in this study because of its flexibility and versatility, including smoothing curves, functional clustering, FPCA, one-way ANOVA tests and newly implemented deep learning methods [51,52]. Nevertheless, we are interested in exploring the application of GP in the analysis of temporal perturbational data in the future.

CellDrift was implemented in two cases in the paper, including immune transcriptome profiling in infectious diseases and organ development over a continuous timeline. The cost of sample collection and single cell sequencing technology is still one of the major obstacles to collecting more longitudinal single cell data. Yet we are beginning to see more large-scale longitudinal single cell experiments due to the popularity of single cell sequencing technology and the progress of organoid research [13]. We established the effective performance of CellDrift in the identification of temporal patterns of gene perturbation effects. Additionally, we can more confidently generate hypotheses for perturbation responses by incorporating gene enrichment analysis. Furthermore, we can associate temporal changes with other clinical events of patients, and apply machine learning methods, such as k-nearest neighbors (KNN), to predict the possibility of certain clinical events of patients before they happen. Notably, other applications of functional data analysis, such as extrapolation and kernel regression, can greatly enhance our ability to evaluate temporal perturbation effects.

There are several important areas that CellDrift and this evaluation do not address. First, we didn’t establish the effectiveness of CellDrift in studies with small numbers (e.g. < 5) of time points. Datasets from many such single-cell studies have been archived and continue to be generated since it is still a relatively costly technology compared with traditional approaches, such as bulk RNA-seq, in large-scale experiments. Additionally, we did not include time as a covariate in the generalized linear model. Instead, the contrast coefficient information was combined from GLM runs of separate time points, which might result in lower statistical power or increased probability of a type 1 error, as well as making the CellDrift procedure cumbersome. This may be an area for future improvement. Additionally, we did not introduce covariance between genes, which would reduce the power of detecting gene correlations of perturbation effects.

## Author’s contributions

Conceptualization K.J, N.S., B.A..; Methodology, K.J., R.S., D.S. and S.P.; Investigation, K.J., G.L., D.S.; Writing, K.J., D.S., G.L., R.S., B.A.; Editing, D.S., G.L., R.S., B.A; Funding Acquisition, B.A.; Resources and Data Curation, K.J.; R.S.; Visualization, K.J.; Supervision, B.A.

## Acknowledgement

We express our appreciation to Dr. Kieran R Campbell and Dr. Catherine Blish for their valuable suggestions on the method design. We appreciate suggestions from Dr. Emrah Gecili for the manuscript revision.

## Data Availability

The source code and Python package are freely available at https://github.com/KANG-BIOINFO/CellDrift. The analysis performed in this paper can be found at: https://github.com/KANG-BIOINFO/CellDrift_analysis.

## Funding

This work was supported by National Institutes of Health LungMap (HL148865), Digestive Health Center (DK078392), National Cooperative Reprogrammed Cell Research Program (MH104172) and Center of Excellence in Molecular Hematology (DK126108).

## Notes

https://github.com/KANG-BIOINFO/CellDrift

https://github.com/KANG-BIOINFO/CellDrift_analysis

### Competing Interest Statement

The authors have declared no competing interest.

