## Supplementary Figures for "CellDrift: Inferring Perturbation Responses in Temporally-Sampled Single Cell Data"

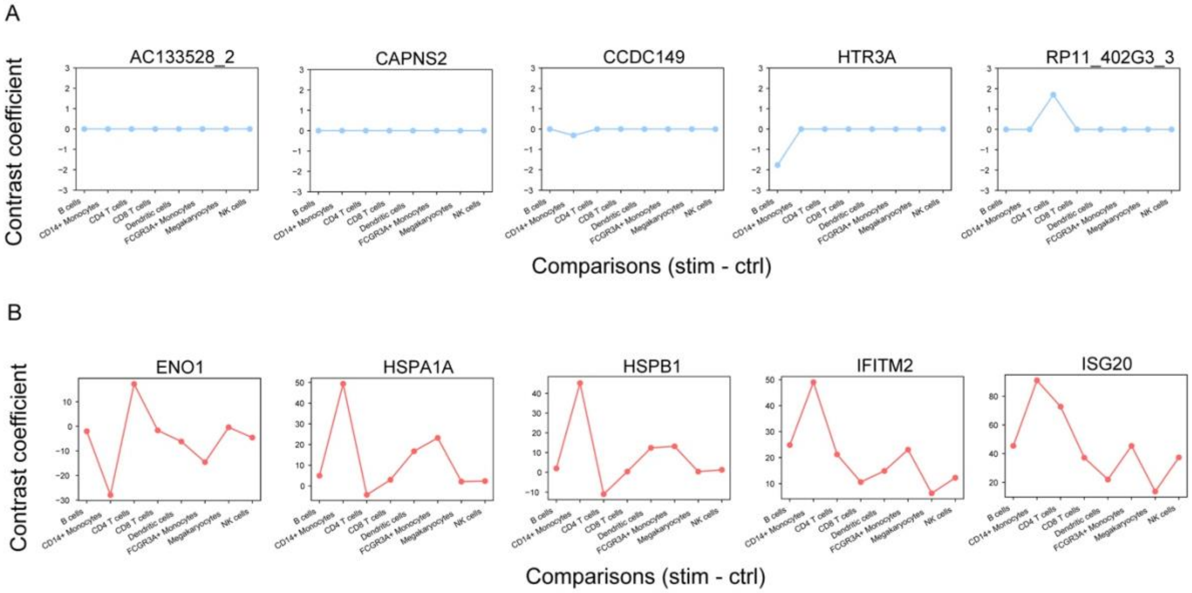


Figure S1. Contrast coefficients of Likelihood Ratio Test (LRT)-prioritized genes. Contrast coefficients of 5 genes with the least (A) and most (B) significant LRT results in the model selection when applying CellDrift in the IFN-stimulated PBMC single-cell data. Genes in (A) displayed minor heterogeneity of perturbation responses (contrast coefficients) across cell types, while genes in (B) displayed great heterogeneity.

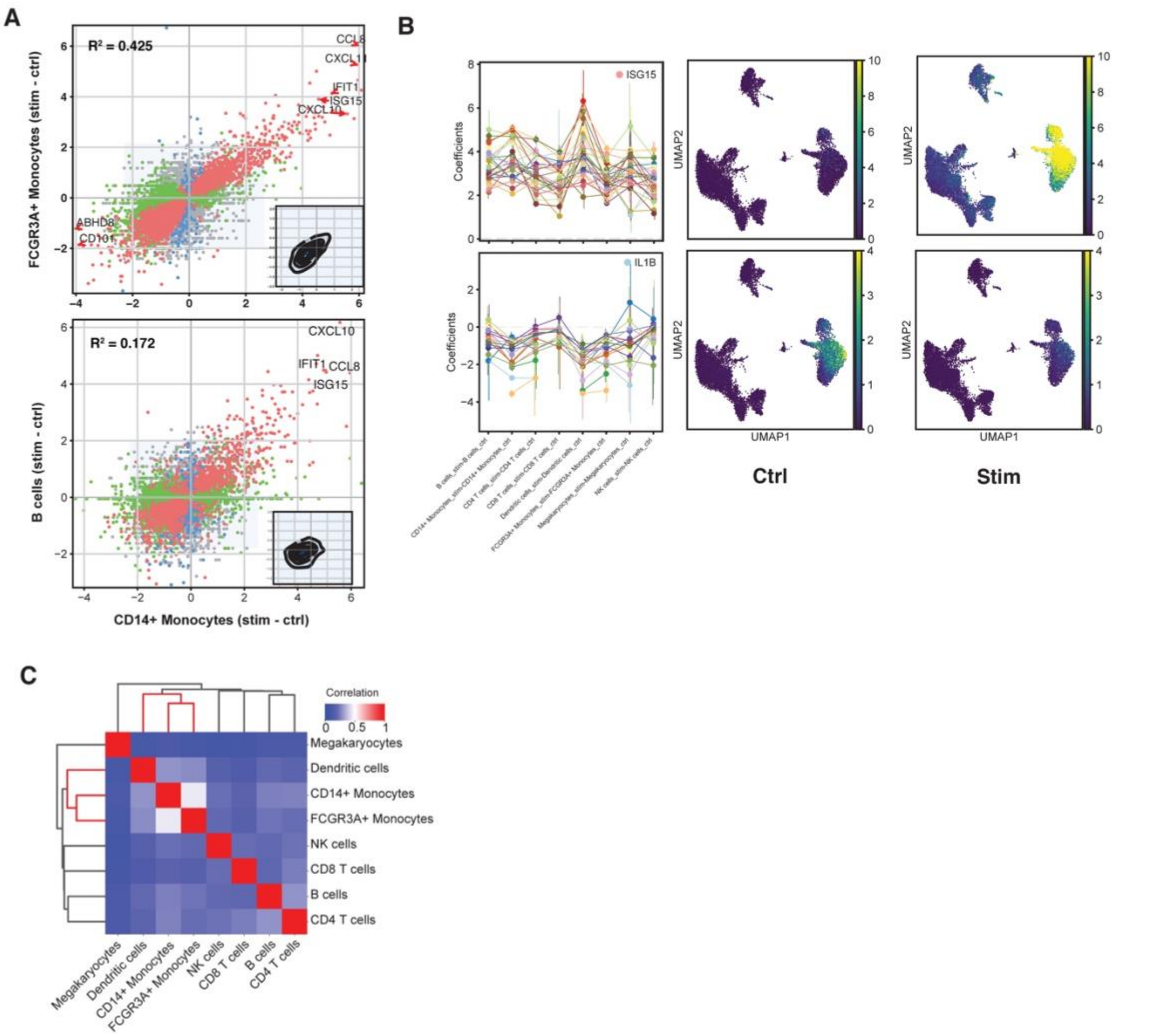


Figure S2. Application of generalized linear model of CellDrift in interferon-stimulated PBMC single cell data. (A) Scatter plots for contrastive coefficients in CD14+ Monocytes versus FCGR3A+ Monocytes (top) and B cells (bottom). Genes that achieve significant levels in both cell types, only one cell type and zero cell type are colored in red, green (or blue) and gray, respectively. $R^2$ scores were derived from a fitted linear model. Similar cell types present closer perturbation responses (top) than dissimilar cell types (bottom). (B) Similar patterns of gene ISG15 (top) and IL1B (bottom) across cell types. The accumulated expression levels of all genes in each pattern are shown in control and interferon-stimulated condition. (C) Pearson correlation matrix of contrast coefficients of all genes across multiple cell types.

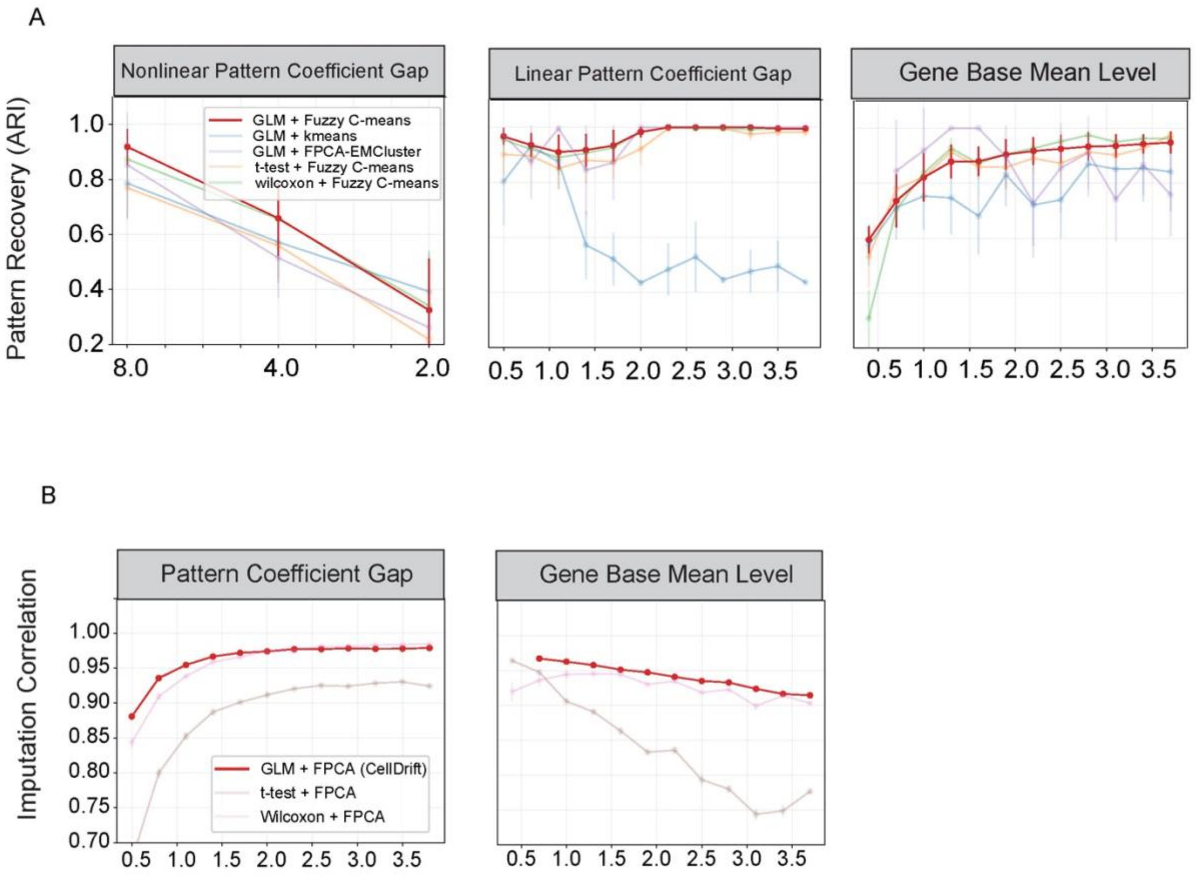


Figure S3. Benchmark of performance of temporal pattern identification and imputation. (A) Performance evaluation of temporal pattern recovery in CellDrift strategy (GLM + fuzzy C-means) versus other strategies upon varying parameters, including gaps of coefficients for non-linear temporal patterns (Left) and linear patterns (Middle), as well as gene base mean level (Right), which represents the sequencing depth of the data. (B) Performance evaluation of Pearson correlation between imputed coefficients and real simulated coefficients across varying pattern coefficient gaps and gene base mean levels.

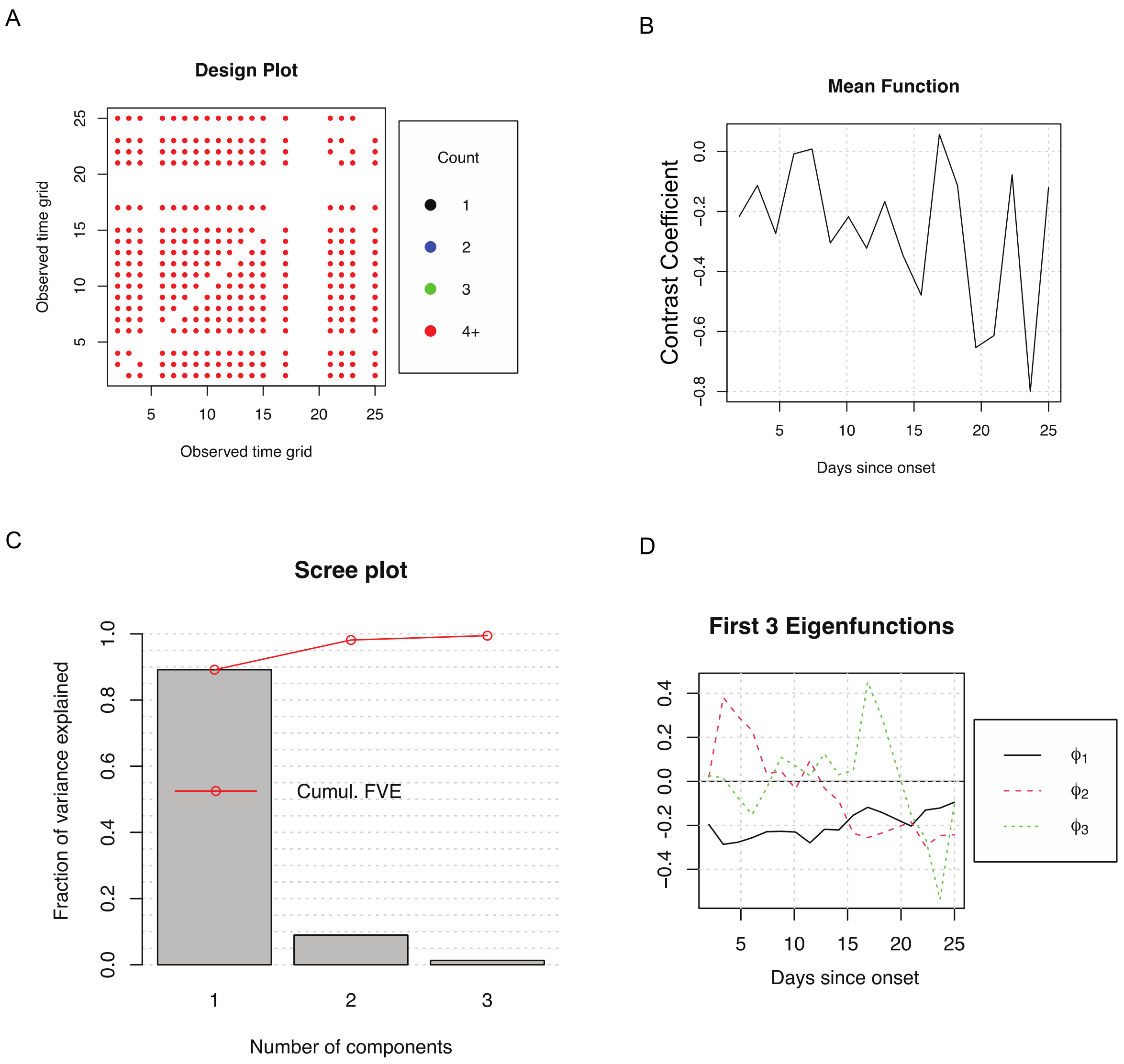


Figure S4. Basic information of FPCA fitting for genes in severe COVID-19 classical Monocytes. (A) The distribution of the number of samples in each time point for input samples. (B) Average contrast coefficients for all input genes across time points. (C) Top three functional principal components explain more than 99% of functional variance. (D) Top three eigenfunctions ($\phi_1, \phi_2, \phi_3$) and their scores across time points.

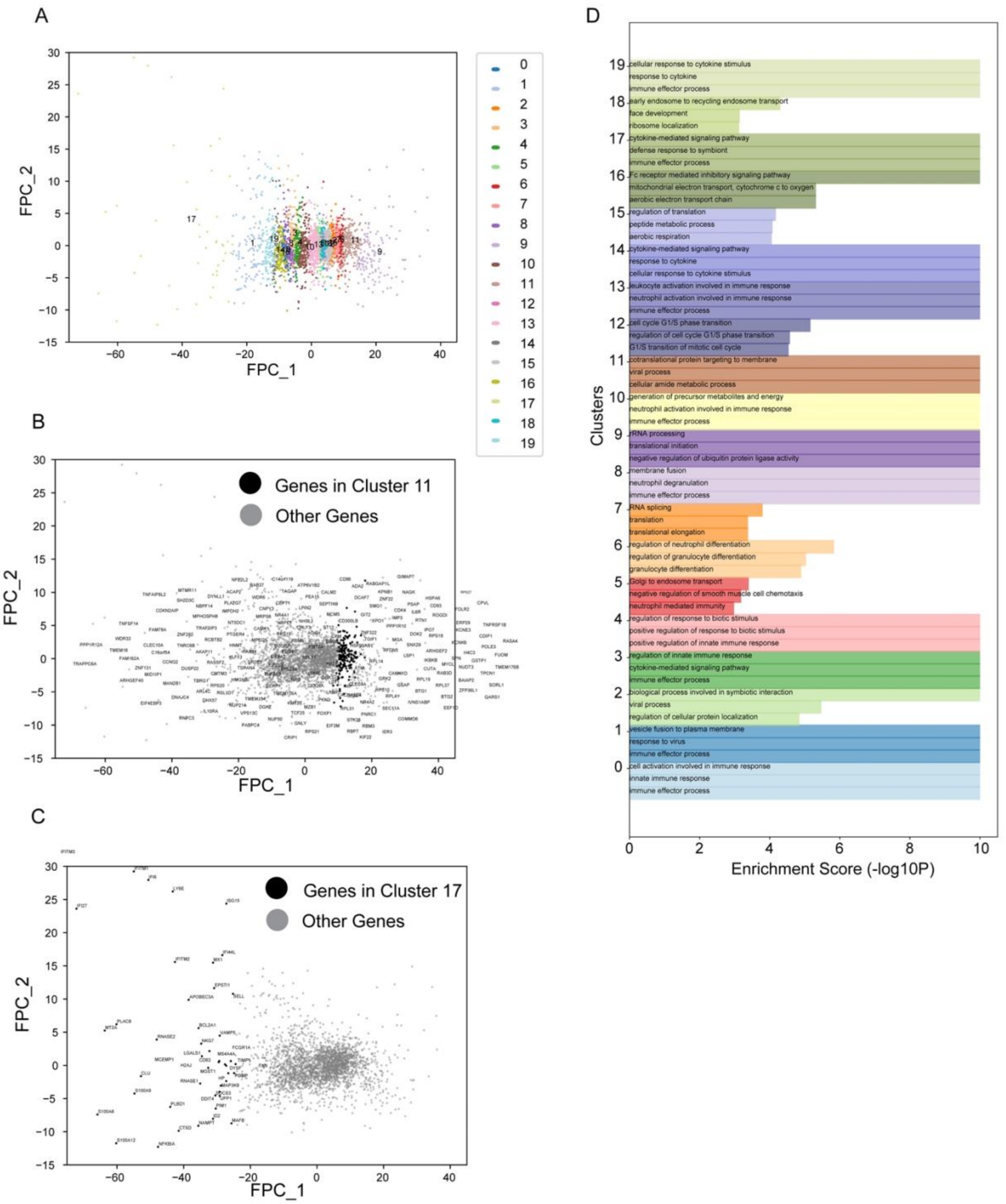


Figure S5. Characteristics of temporal clusters for classical monocytes in severe COVID-19. (A) Distribution of 20 temporal clusters on 2D functional PC dimensions. (B-C) Highlight of genes in cluster 11 (B) and 17 (C) in an enlarged visualization of (A). (D) Gene enrichment analysis for each temporal cluster. Biological Process from Gene Ontology was used to annotate gene functions. Enrichment scores were defined as $-log_{10}FDR - adjusted\ p\ values$ of enrichment significance and trimmed at 10.

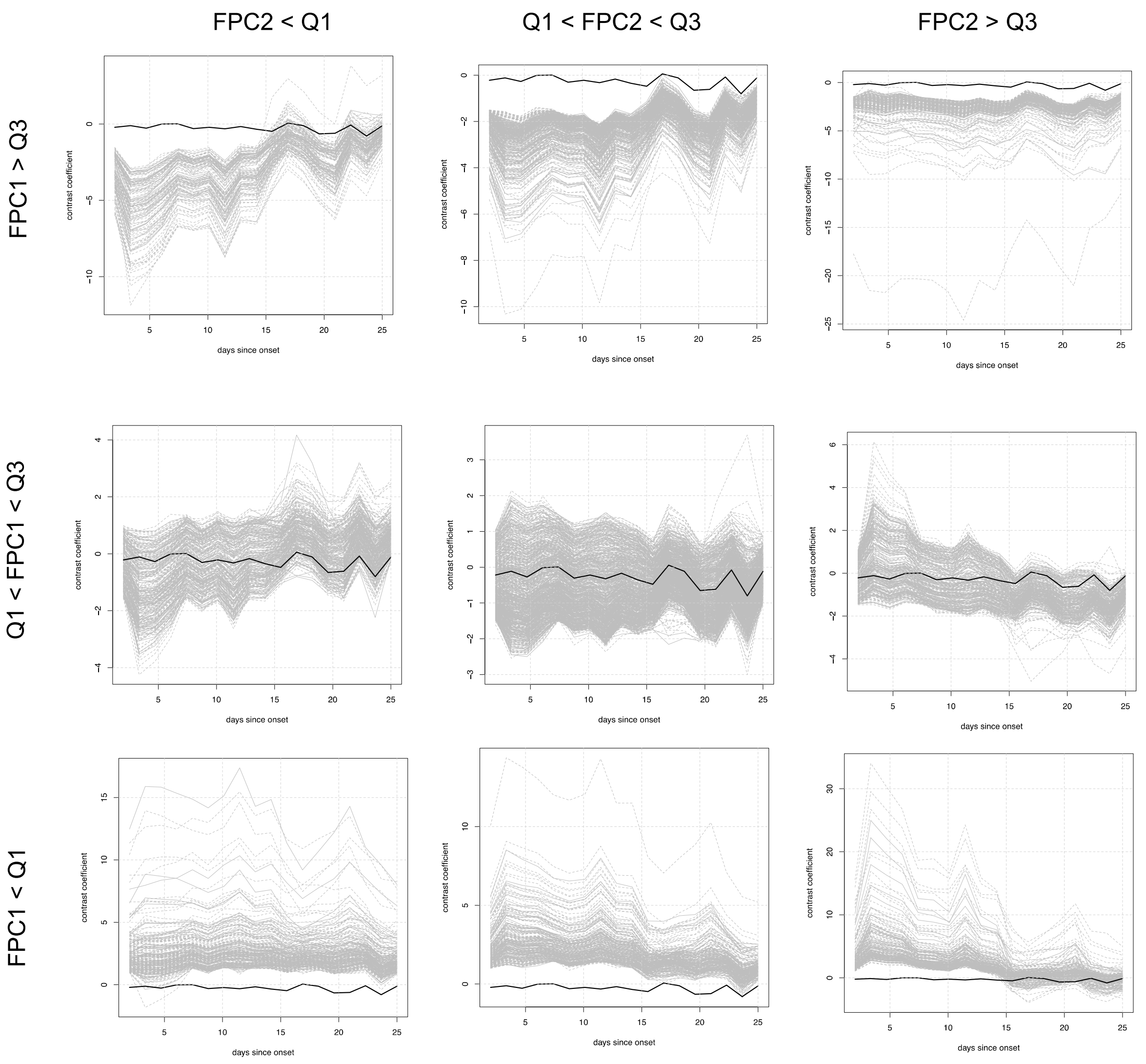


Figure S6. Stratification of genes with the scores of first to functional principal components. Functional principal components scores of genes were stratified into 3 sections, including < Q1 (bottom 25%), Q1 - Q3 (25% - 75%) and >Q3 (top 25%). Average contrast coefficient for all genes is shown in a black curve.

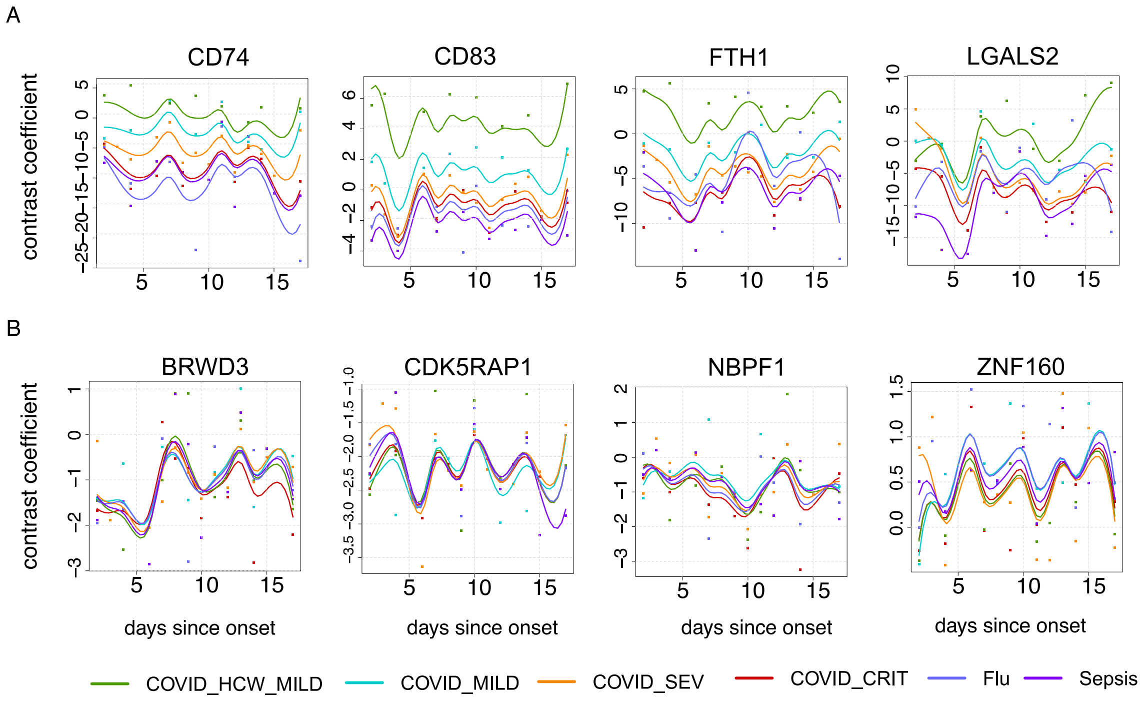


Figure S7. One-way ANOVA results for the disease condition comparison. (A) Four genes from genes with highest ANOVA statistical scores by comparing mild conditions (HCW mild and mild COVID-19) with severe conditions (severe and critical COVID-19, severe influenza and sepsis), where mild conditions have higher average temporal curve than severe conditions. (B) Four genes from genes with lowest ANOVA statistical scores by comparing mild and severe conditions, representing genes with little temporal variation for those two disease conditions.

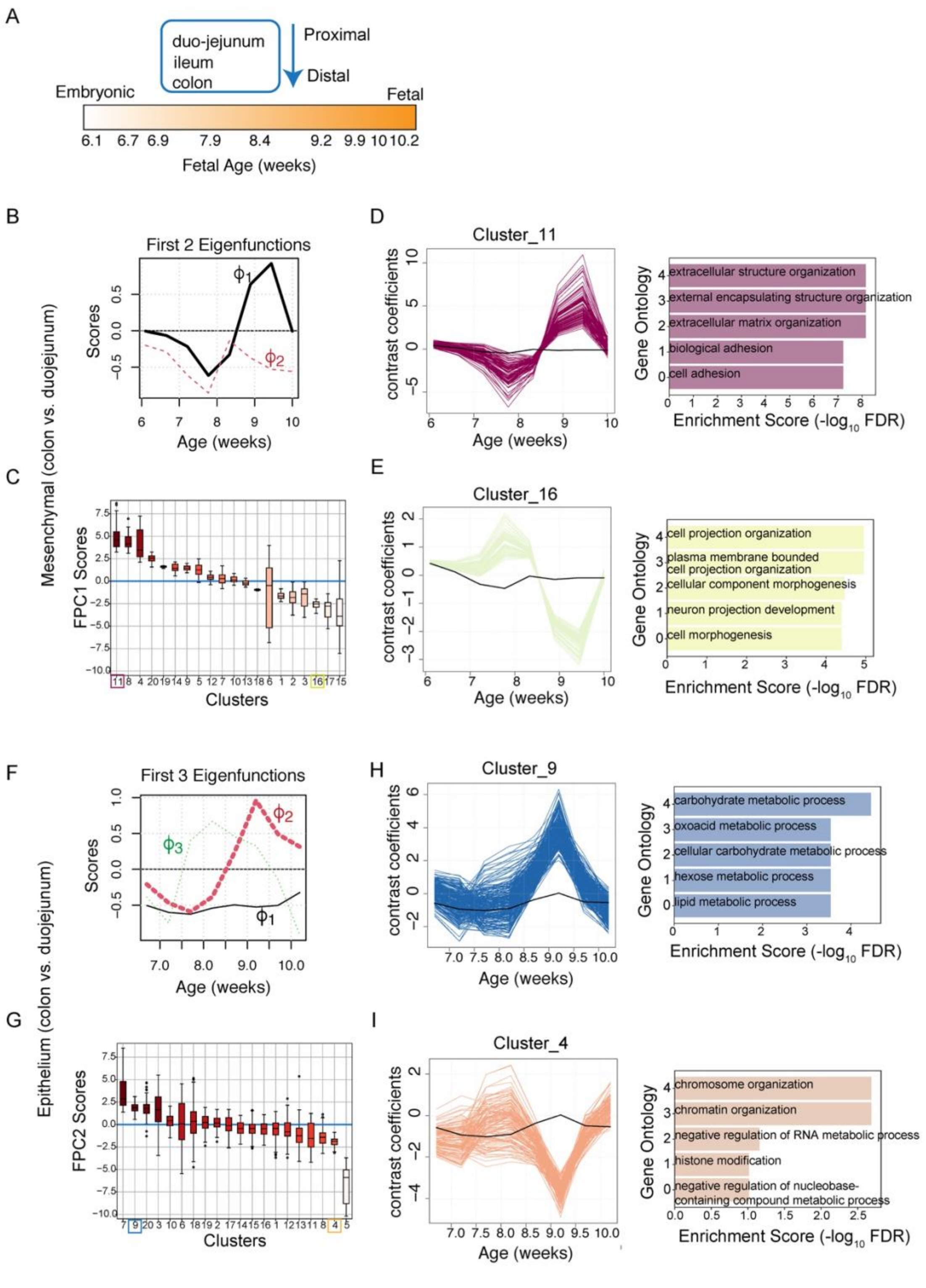


Figure S8. Differential temporal trajectories of gut development. (A) Overview of fetal gut cell atlas. The study contains single cell sequencing data for 3 compartments in gut development, including duo-jejunum, ileum and colon, whose transcriptomics profiles were retrieved from week 6 to week 11. (B) Top 2 FPCA eigenfunctions and their scores over time for functional data of comparisons of colon and duo-jejunum in mesenchymal cells. First eigenfunction $\phi_1$ presents a strong temporal pattern as is highlighted. (C) Functional principal component1 (FPC1) scores were computed for genes in each cluster, displayed in decreasing order. High positive scores indicates genes in the cluster may have similar temporal patterns as $\phi_1$, while large negative scores indicate opposite temporal patterns as $\phi_1$. Clusters 11 and 16 are highlighted for the following analysis. (D-E) FPCA smoothed curves for genes in cluster 11 (D) and 16 (E) and their gene enrichment results. (F) Top 3 FPCA eigenfunctions and their scores over time for functional data of comparisons of colon and duo-jejunum in epithelium cells. Second eigenfunction $\phi_2$ presents a strong temporal pattern as is highlighted. (G) Functional principal component 2 (FPC2) scores were computed for genes in each cluster, which was ranked in a decreasing order. Clusters 4 and 9 are highlighted for the following analysis. (H-I) FPCA smoothed curves for genes in cluster 9 (H) and 4 (I) and their gene enrichment results.

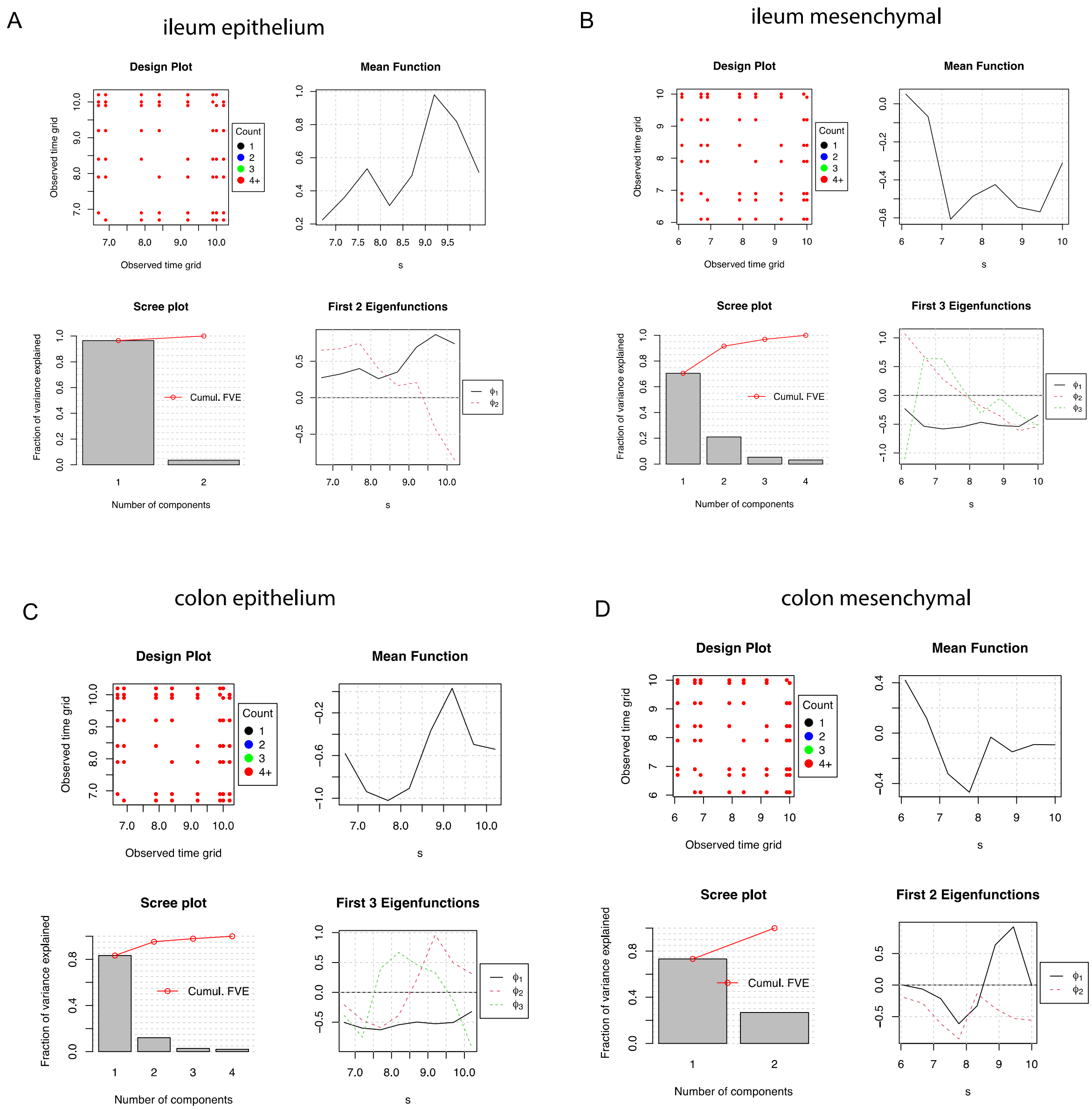


Figure S9. Basic information of FPCA for four comparisons, including ileum vs. duo-jejunum in epithelium (A); ileum vs. duo-jejunum in mesenchymal (B); colon vs. duo-jejunum in epithelium (C) and colon vs. duo-jejunum in mesenchymal (D). More detailed explanations can be found in Figure S5.

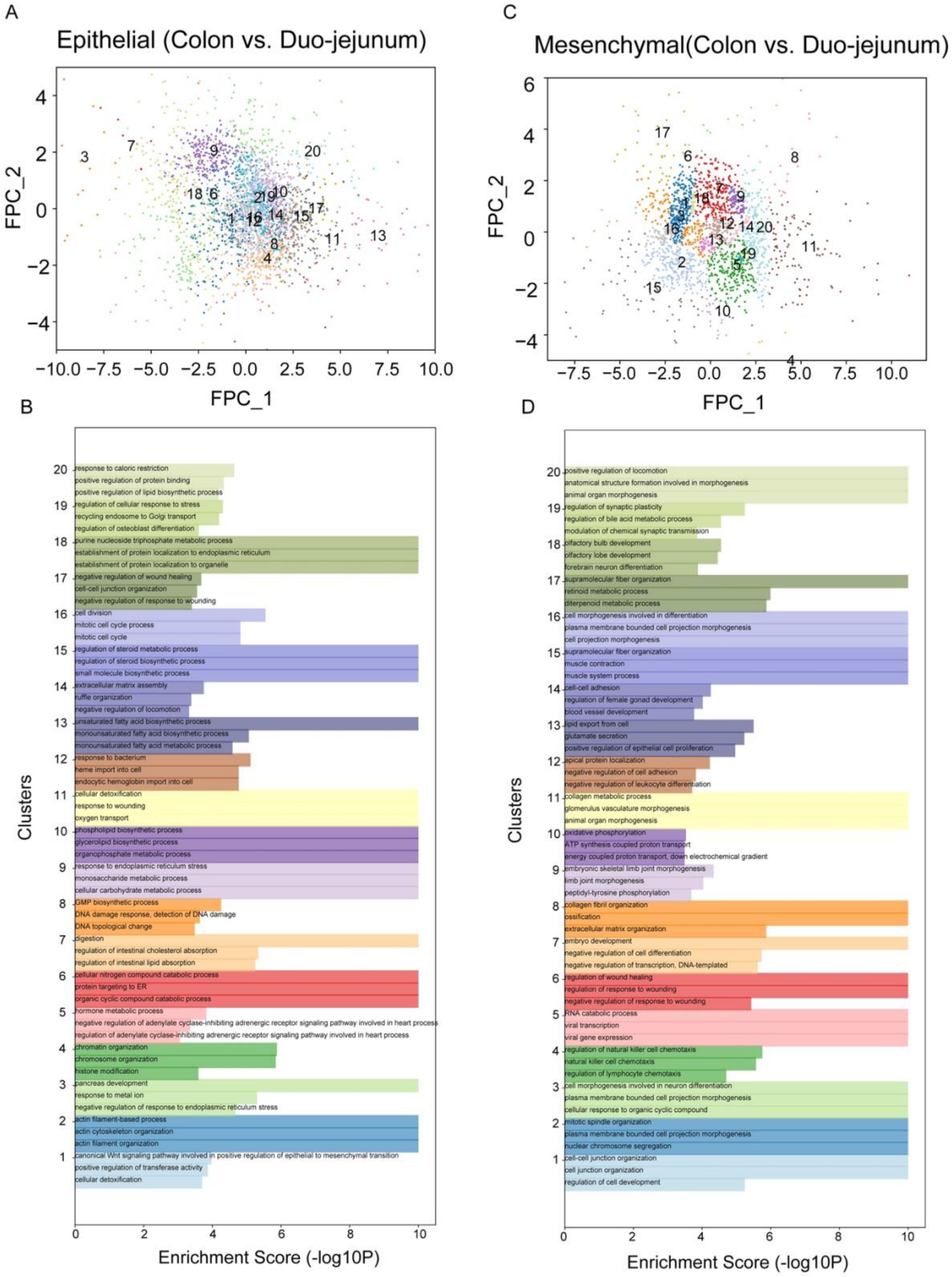
A
Epithelial (Colon vs. Duo-jejunum)
FPC_2
FPC_1
C
Mesenchymal(Colon vs. Duo-jejunum)
FPC_2
FPC_1
B
Clusters
Enrichment Score (-log10P)
response to caloric restriction
positive regulation of protein binding
positive regulation of lipid biosynthetic process
regulation of cellular response to stress
recycling endosome to Golgi transport
regulation of osteoblast differentiation
purine nucleoside triphosphate metabolic process
establishment of protein localization to endoplasmic reticulum
establishment of protein localization to organelle
negative regulation of wound healing
cell-cell junction organization
negative regulation of response to wounding
cell division
mitotic cell cycle process
mitotic cell cycle
regulation of steroid metabolic process
regulation of steroid biosynthetic process
small molecule biosynthetic process
extracellular matrix assembly
ruffle organization
negative regulation of locomotion
unsaturated fatty acid biosynthetic process
monounsaturated fatty acid biosynthetic process
monounsaturated fatty acid metabolic process
response to bacterium
heme import into cell
endocytic hemoglobin import into cell
cellular detoxification
response to wounding
oxygen transport
phospholipid biosynthetic process
glycerolipid biosynthetic process
organophosphate metabolic process
response to endoplasmic reticulum stress
monosaccharide metabolic process
cellular carbohydrate metabolic process
GMP biosynthetic process
DNA damage response, detection of DNA damage
DNA topological change
digestion
regulation of intestinal cholesterol absorption
regulation of intestinal lipid absorption
cellular nitrogen compound catabolic process
protein targeting to ER
organic cyclic compound catabolic process
hormone metabolic process
negative regulation of adenylate cyclase-inhibiting adrenergic receptor signaling pathway involved in heart process
regulation of adenylate cyclase-inhibiting adrenergic receptor signaling pathway involved in heart process
chromatin organization
chromosome organization
histone modification
pancreas development
response to metal ion
negative regulation of response to endoplasmic reticulum stress
actin filament-based process
actin cytoskeleton organization
actin filament organization
canonical Wnt signaling pathway involved in positive regulation of epithelial to mesenchymal transition
positive regulation of transferase activity
cellular detoxification
D
Clusters
Enrichment Score (-log10P)
positive regulation of locomotion
anatomical structure formation involved in morphogenesis
animal organ morphogenesis
regulation of synaptic plasticity
regulation of bile acid metabolic process
modulation of chemical synaptic transmission
olfactory bulb development
olfactory lobe development
forebrain neuron differentiation
supramolecular fiber organization
retinoid metabolic process
diterpenoid metabolic process
cell morphogenesis involved in differentiation
plasma membrane bounded cell projection morphogenesis
cell projection morphogenesis
supramolecular fiber organization
muscle contraction
muscle system process
cell-cell adhesion
regulation of female gonad development
blood vessel development
lipid export from cell
glutamate secretion
positive regulation of epithelial cell proliferation
apical protein localization
negative regulation of cell adhesion
negative regulation of leukocyte differentiation
collagen metabolic process
glomerulus vasculature morphogenesis
animal organ morphogenesis
oxidative phosphorylation
ATP synthesis coupled proton transport
energy coupled proton transport, down electrochemical gradient
embryonic skeletal limb joint morphogenesis
limb joint morphogenesis
peptidyl-tyrosine phosphorylation
collagen fibril organization
ossification
extracellular matrix organization
embryo development
negative regulation of cell differentiation
negative regulation of transcription, DNA-templated
regulation of wound healing
regulation of response to wounding
negative regulation of response to wounding
RNA catabolic process
viral transcription
viral gene expression
regulation of natural killer cell chemotaxis
natural killer cell chemotaxis
regulation of lymphocyte chemotaxis
cell morphogenesis involved in neuron differentiation
plasma membrane bounded cell projection morphogenesis
cellular response to organic cyclic compound
mitotic spindle organization
plasma membrane bounded cell projection morphogenesis
nuclear chromosome segregation
cell-cell junction organization
cell junction organization
regulation of cell development

Figure S10. Characteristics of temporal clusters in gut development. (A, C) Distributions of 20 temporal clusters on 2D FPC dimensions for colon vs. duo-jejunum in epithelium (A) and mesenchymal cells (C). (B, D) Gene enrichment results for 20 clusters in (A) and (C).
