## Supplementary Methods for "CellDrift: Inferring Perturbation Responses in Temporally-Sampled Single Cell Data"

**Dataset Description**

(1) Interferon-stimulated PBMC [[1]](https://paperpile.com/c/xSeiVd/cdQn): The dataset was downloaded from GEO accession GSE96583. Before sequencing, samples were thawed for 6 hours with or without IFN-beta stimulation. Based on the original annotations, doublets were removed from the input cell.

(2) COVID-19 Atlas (COMBAT) [[2]](https://paperpile.com/c/xSeiVd/4Gn5): This is a large cohort of single-cell CITE-seq study of

PBMC cells from control donors, COVID-19, influenza and sepsis patients. 116 hospitalized COVID-19 patients were recruited from a single site, including mild, mild (healthcare workers), severe and critical COVID-19 patients. Disease progression days (0~25 days since onset) were recorded for participants in the study. We extracted the six most common cell types in the study for the CellDrift analysis.

(3) COVID-19 Atlas (Ren et al.) [[3]](https://paperpile.com/c/xSeiVd/Y1yU): This is a large COVID-19 cohort of single-cell data with 284 samples from 196 COVID-19 patients. 1.46 million cells were collected from multiple tissues, including bronchoalveolar lavage fluid (BALF), sputum, peripheral blood mononuclear cells (PBMC) and pleural fluid mononuclear cells (PFMCs). Control donors, mild/moderate and severe/critical COVID-19 patients were recruited. The data was downloaded from GEO accession GSE158055 and used as validation data in our study. We extracted PBMC cells from control donors and patients in progression (patients during convalescence were not included in the analysis) for downstream analysis. B cells, CD4+ T cells, CD8+ T cells and Monocytes were included in CellDrift analysis.

(4) Fetal Gut Atlas [[4]](https://paperpile.com/c/xSeiVd/21EY): This data contains single cell transcriptomics from human embryos with a post-conceptional age ranging from 6 to 10 weeks. Dissected tissues were divided into proximal small bowel (duodenum and jejunum, or duo-jejunum), distal small bowel (ileum), and large bowel (colon). Four lineages of cells, including epithelium, immune, mesenchymal and vasculature, were included in the data. We used epithelium and mesenchymal cells for downstream analysis since there are enough cells in these two lineages for the CellDrift input. The data was downloaded from Gut Cell Atlas: [https://www.gutcellatlas.org](https://www.gutcellatlas.org/fetal).

**Differential Expression Methods in Benchmark**

We benchmarked four differential expression methods in the study, including t-test, t-test (overdispersion), wilcoxon test and MAST, which are popular DE methods in single cell analysis. T-test, t-test (overdispersion) and wilcoxon test were implemented in Scanpy [[6]](https://paperpile.com/c/xSeiVd/8Jsb) using function $rank\_genes\_groups$. Compared with t-test, t-test (overdispersion) forces the variance of cells in both comparison groups to be the same to mitigate the statistical issues caused by imbalanced cell numbers. It was the default DE method in previous Scanpy versions. MAST is a generalized linear model that adjusts the cellular detection rate and the fraction of genes expressed in a cell in the differential expression analysis [[7]](https://paperpile.com/c/xSeiVd/2GHB). Compared with other methods, it has more flexibility to address the nuisance variation in single cell count data using the GLM framework.

**Functional clustering algorithms in Benchmark**

We benchmarked three popular functional clustering algorithms in our study, including KMeans (KM), Fuzzy C-means (FCM) and FPCA-based EMCluster.

KMeans and Fuzzy C-means were implemented in scikit-fda [[8]](https://paperpile.com/c/xSeiVd/4W1g), where the distance for pairwise observations $f$ and $g$ were measured by $lp\_distance$:

$d(f,g) = d(g,f) = {|| f- g ||}_{p}$

where ${||\cdot||}_{p}$ denotes the Lp norm. F	or each observation $f$ the Lp norm is defined as:

$||f|| = \left( \int_{D} {||f||}^{p}dx \right)^{\frac{1}{p}}$

where D is the domain over which the functions are defined. Default p is equal to 2.

For KMeans functional clustering, let $X = \{x1, x2, ..., xn \}$ be the functional data to be analyzed, and $V = \{v1, v2, .., vc\}$ be the set of center of clusters in X dataset, where $n$ is the number of observations and $c$ is the number of clusters. KMeans iteratively computes cluster centroids to minimize the square error function:

$J_{KM}(X; V) = \sum_{i = 1}^{C} \sum_{j = 1}^{n} D_{ij}^{2}$

where $D_{ij}^{2}$ is the squared distance measured by p-norm:

$$D_{ij}^{2} = ||x_{ij} - v_{i}|| = \int_{I} {({|x_{ij} - v_{i}|}^{p}dx)}^{\frac{1}{p}}$$

Different from KMean clustering, Fuzzy C-means clustering aims to minimize the weighted square errors instead of square errors:

$J_{FCM}(X; U, V) = \sum_{i = 1}^{C} \sum_{j = 1}^{n} u_{ij}^{f}D_{ij}^{2}$

where U is a fuzzy partition matrix that represent membership values of observations to each cluster:

$u_{ij}^{f} = \left[ \sum_{k =1}^{c} {(D_{ij} / D_{kj})}^{\frac{2}{f-1}} \right]^{-1}$

Cluster centroids $V$ are updated by the following formula:

$v_{i} = \frac{\sum_{j=1}^{n} u_{ij}^{f}x_{j}}{\sum_{j=1}^{n} u_{ij}^{f}}, 1\leq i \leq c$

FPCA-based EMCluster was implemented using function $FClust$ from R package $fdapace$. In $fdapace$, functional principal component scores are first calculated using FPCA in $fdapace$ and then clustering is performed using EMCluster algorithm. EMCluster assumes a finite mixture Gaussian distribution with unstructured dispersion and applies an EM algorithm for model-based clustering. More details of it can be found in R package $EMCluster$.

**Functional Principal Component Analysis**

Functional principal component analysis (FPCA) was implemented using the R package fdapace, where Principal Analysis by Conditional Estimation (PACE) was performed to retrieve functional PCs [[5]](https://paperpile.com/c/xSeiVd/Whuo). Here is a brief introduction of PACE procedure in the package (revised based on tutorial of fdapace):

1. Smoothed mean $\hat{\mu}$ is calculated using local linear smoothing and all the available readings are aggregated together.

2. Raw covariance for each curve is calculated separately and then all raw covariances are aggregated to generate the sample raw covariance.

3. Smoothed covariance is estimated using the off-diagonal elements of the sample raw covariance.

4. Eigenanalysis is performed on the smoothed covariance to obtain the estimated eigenfunctions $\hat{\phi}$ and eigenvalues $\hat{\lambda}$. The smoothed covariance is then projected on a positive semi-definite surface.

5. Conditional Expectation (PACE step) is used to estimate the corresponding scores $\hat{\xi}$:

$\hat{\xi}_{ik} = \hat{E}[\hat{\xi}_{ik} | Y_{i}] = {\hat{\lambda}_{k}\hat{\phi}_{ik}^{T}\sum_{Y_{i}}^{-1} (Y_{i} - \hat{\mu}_{i})}$

**One way ANOVA functional test**

One-way function ANOVA was implemented using the $oneway\_anova$ function in the package scikit-fda. It is used to analyze whether there is a difference among means of the functional data.

Let ${\{X_{i}\}}_{i=1}^{k}$ be $k$ independent samples and each one with $n_{i}$ trajectories; let $E(X_{i}) = m_{i}(t)$. The null hypothesis is:

$$H_{0}: m_{1}(t) = m_{2}(t) = {... = m}_{k}(t)$$

$oneway\_anova$ calculates a statistic that measures the variability between groups of samples:

$$V_{n} = \sum_{i<j}^{k} w_{i}{||f_{i} - f_{j}||}^{2}$$

where ${\{f_{i}\}}_{i = 1}^{k}$is a set of samples in the functional data; ${\{w_{j}\}}_{j = 1}^{k}$ is a set of weights, where $w_{i}$is related to the sample $f_{i}$ for $i =1,...,k$.

Under the null hypothesis this statistic $V_{n}$ is asymptotically equivalent to $v\_asymptotic\_stat()$, where each sample is replaced by a gaussian process with zero mean and the same covariance as the original curve. More details of the procedure can be found in the original paper [[9]](https://paperpile.com/c/xSeiVd/ML5p).
